# Differential lipid dynamics in stocked and wild juvenile lake trout

**DOI:** 10.1101/718692

**Authors:** Madelyn G. Sorrentino, Taylor R. Stewart, J. Ellen Marsden, Jason D. Stockwell

## Abstract

After more than 40 years of stocking, lake trout (*Salvelinus namaycush*) in Lake Champlain have started to exhibit strong, natural recruitment. The abrupt surge in recruitment suggests a change in limiting factors such as prey availability or overwinter survival. The distribution of juvenile wild lake trout varies in relative abundance among regions of Lake Champlain. The differences suggest the prey base, or foraging success, may vary geographically within the lake. Stocked and wild lake trout may differ in their ability to use resources and in overwinter survival. One metric that can indicate differences in resources across regions is lake trout lipid content, which reflects the quality of available food and serves as an important energy reserve for overwinter survival. We quantified total lipid content of stocked and wild juvenile lake trout across spatial (lake regions) and temporal (seasonal) scales. No spatial differences in lipid content were apparent. Wild fish had greater lipid content than stocked fish. Seasonally, stocked fish showed a continuous drop in lipid content from pre-winter levels at stocking to the following autumn. Wild fish showed a cyclical summer increase in lipids following winter depletion, which plateaued by autumn. The high lipid content of hatchery lake trout may be necessary as they acclimate to foraging in the wild. Hatcheries would benefit from evaluating whether post-stocking survival could be improved by altering feeding or rearing regimes.

## Introduction

Lake trout *(Salvelinus namaycush)* was extirpated from Lake Champlain by 1900 (Plosila and Anderson, 1985). Restoration efforts began in 1972 with an intensive stocking program to reestablish a self-sustaining population and a recreational fishery (Marsden et al., 2010; Marsden and Langdon, 2012). Successful spawning and fry emergence were documented at several sites starting in 2000 but sustained natural recruitment did not begin until 2012, four decades after the stocking program commenced (Marsden et al., 2018). Recent natural recruitment may be due to a change in limiting factors such as food quality or quantity. For example, the Lake Champlain prey base was diversified in 2003 by the invasion of alewife (*Alosa pseudoharengus*), a known diet item of juvenile lake trout (Marsden et al. unpublished data; Madenjian et al., 2006). Winter is a period of high mortality for juvenile fishes when the risks of starvation, thermal stress, and predation is high (Hjort, 1914; Hurst, 2007). Increased prey availability, milder winter conditions, or other factors could help juvenile lake trout survive the winter critical period.

Juvenile surveys indicate that relative abundance of stocked and wild lake trout varies across regions of Lake Champlain. Annual stocking occurs at two highly productive spawning sites, Whallon Bay in the southern Main Lake and Gordon Landing in the northern Main Lake (Ellrott and Marsden, 2004). However, the highest proportion and relative abundance (catch-per-unit-effort, CPUE) of wild fish has been consistently found in the central Main Lake (Marsden et al., 2018; Wilkins et al. in review). The difference in expected versus observed distributions suggest that prey resources may be asymmetrically distributed across the lake, unknown but productive spawning sites exist in the central Main Lake, or both.

An indirect measure of high foraging success, or decreased winter stress, in fishes is lipid storage. Lipid content in juvenile lake trout could provide insight into the recent surge in natural recruitment because of its roles in fish health – lipids serve as energy resources and help fish to cope with environmental stressors (Adams, 1999; Tocher, 2003). In particular, lipids are used for basic maintenance and other metabolic needs during winter, when prey availability is presumably low and typically reduced by the end of the season (Adams, 1999; MacKinnon, 1972; Rikardsen and Elliott, 2000). For example, juvenile rainbow trout (*Oncorhynchus mykiss*) and juvenile Atlantic salmon (*Salmo salar*) exhibited depleted lipid reserves (60-90% and 34-57% depletion, respectively) over winter (Biro et al., 2004; Naesie et al., 2006). Additionally, the health of fish can often be predicted by lipid content; fish with low growth and condition factor have correspondingly low lipid content (Amara et al., 2007). Accordingly, total lipid content provides an assessment of the energy status of a fish (Naesie et al., 2006; Trudel et al., 2005), and may indicate how well fish are prepared to survive the winter and how they respond to winter depletion of energy reserves. Differences in lipid content may help explain why lake trout in Lake Champlain are exhibiting natural recruitment and how different areas of the lake might support the growth of juvenile wild fish. Variation in lipid content between stocked and wild juvenile fish could also reveal differences in the abilities of wild and stocked fish to survive stressors such as the winter season.

We hypothesized that total lipid content of wild juvenile lake trout would be greatest in the central Main Lake where wild recruits are most abundant (Marsden et al., 2018; Wilkins et al. in review), and would be highest in the summer when the prey base is most abundant. We also hypothesized that recently stocked lake trout would have a higher lipid content than wild juveniles because hatchery fish are typically fed a highly nutritious diet under ideal conditions prior to their release. However, post-release stress and adaptation to a wild-caught diet could result in a substantial reduction in lipid content. To test our hypotheses, we measured total lipid content of stocked and wild juvenile lake trout (ages 0-3) in Lake Champlain from three areas of the Main Lake basin during three seasons, and lipid content of age-0 hatchery lake trout prior to stocking.

## Methods

### Study System

Lake Champlain is situated between New York and Vermont, USA, and Quebec, Canada (Figure 1). The lake is 193 km long, with a maximum width of 20 km. The Main Lake is meso-oligotrophic, with a maximum depth of 122m. Since 1995 lake trout have been primarily stocked at Whallon Bay, Gordon Landing, and Burlington Bay (Figure 1; Marsden et al., 2018).

**Figure 1:**
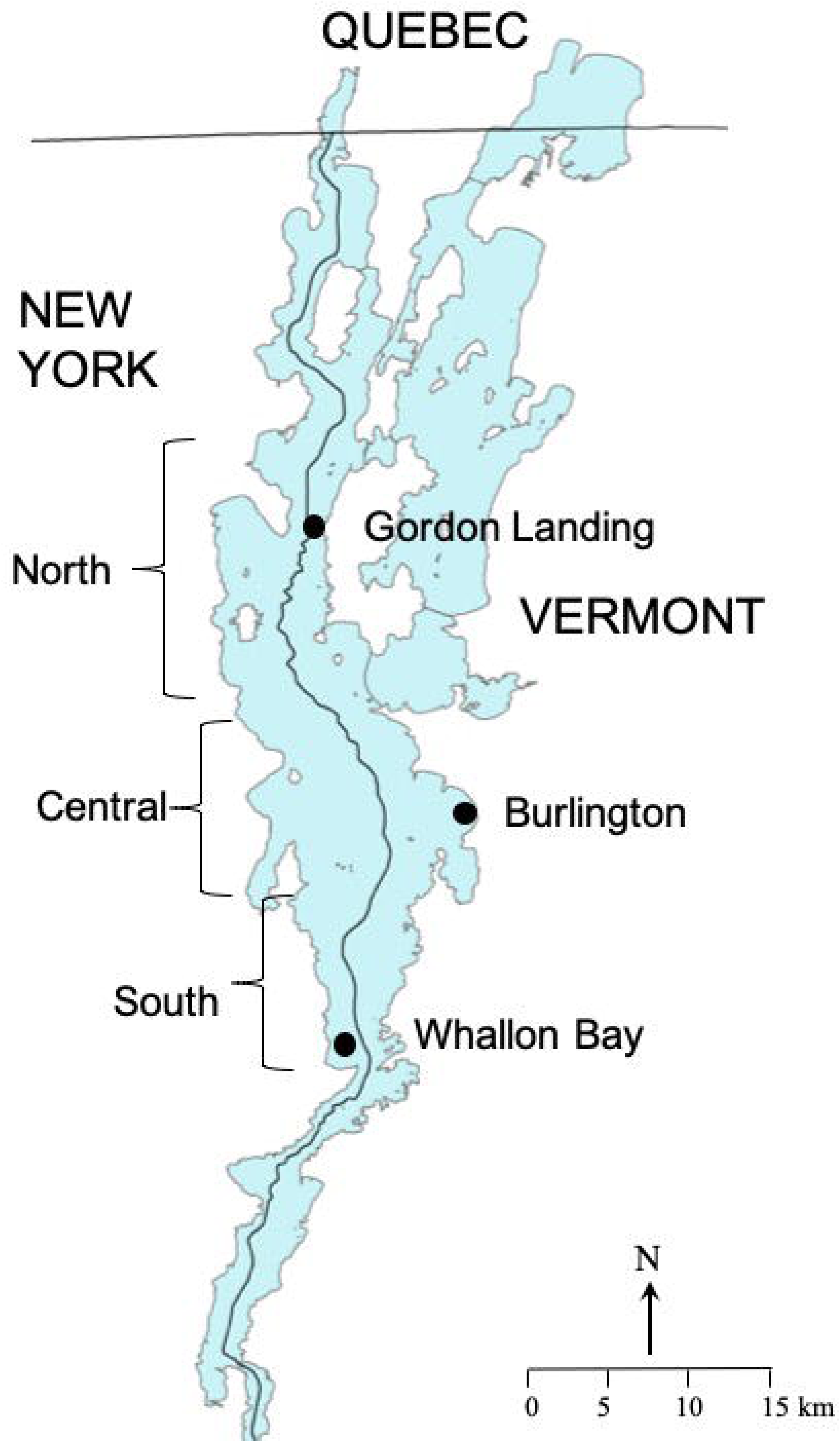
Lake Champlain, showing sampling sites in the Main Lake and two major known lake trout spawning sites at Gordon Landing and Burlington Bay.

### Sample Collection

Fish were sampled at three areas in the Main Lake, near Burlington Bay, Whallon Bay, and Grand Isle (hereafter referred to as the central, south, and north sites) (Figure 1). Sampling efforts for juvenile lake trout have been concentrated at these locations over the past four years, and provided information on variation in relative abundance of stocked and wild lake trout throughout the Main Lake (Marsden et al., 2018).

Sampling was conducted between 8 June and 28 September 2018 to assess potential seasonal changes in lake trout condition. The central site was sampled every 2-3 weeks, and north and south sites were each sampled twice (June and August). We used a three-in-one bottom trawl with an 8-m headrope, 9.3-m footrope with chains, and 1.25-mm stretch cod end liner (Marsden et al., 2018). Trawl tows were taken along-contour at depths from 28 m to 64 m, with the majority of tows concentrated around 40 m, for 10 or 20 min at ∼5.5 km/h. Approximately 30 lake trout were selected from the trawls on each sampling date to represent the range of sizes captured up to 300 mm, and included both stocked and wild fish from each site (i.e., 15 stocked and 15 wild fish). Stocked fish were identified based on presence of a fin clip (Marsden et al. 2018). Fish were immediately frozen on dry ice and stored at −80°C until lipid extraction. A sample of hatchery-reared lake trout (15 fish) was collected from the Ed Weed Fish Culture Station, Grand Isle, VT, on 15 November 2018 to assess lipid content of the lake trout a week prior to release into Lake Champlain. Lake trout are stocked in Lake Champlain at age-0, but are reared to a size equivalent to age-1 wild fish (Marsden et al. 2018).

### Sample Preparation

All lake trout were thawed and measured (total length), weighed, re-assessed for fin clips, and aged based on fin clips and non-overlapping size classes. Fish were dissected and stomach contents removed to avoid any influence of recently consumed prey on the estimate of total lipid content. Each lake trout >150 mm in total length was homogenized in a Ninja BL500 Professional Blender, and a 30-g subsample was removed. Lake trout <150 mm in total length were dried whole. Subsamples and whole small fish were dried to a constant mass at 65°C for 72 hours. Once dry, samples were ground in a mortar and pestle to produce a fine powder.

### Lipid Extractions

Three 1-g (for lake trout >150mm) or 0.5-g (for lake trout <150mm) samples were measured from the dried mass of each fish, and placed into pre-weighed 50-ml conical centrifuge tubes. Samples were analyzed for total lipid content according to a modified version of the Folch et al. (1957) method. Briefly, 10 or 20 ml (depending on sample weight) of a 2:1 chloroform:methanol solution was added to each centrifuge tube. Samples were agitated for 30 seconds using a vortex, and centrifuged for 10 minutes at 3,000 rpm. The lipid-containing supernatant was carefully pipetted off to avoid disturbing the pellet, and the process was repeated a second time. The resulting pellets were then dried for 24 hr at 65°C to ensure evaporation of any remaining chloroform:methanol solution. Samples were weighed again in the centrifuge tubes to estimate the final lipid-free dry mass measurement.

### Data Analysis

Mean percent total lipid content (MPTLC) of the dry fish weight was determined by dividing the pre-extraction weight of each sample by the post-extraction weight and converting to a percent, after subtracting the weight of each centrifuge tube. The percent total lipid content of the three subsamples per fish was averaged together to estimate the MPTLC for each fish. MPTLC was transformed using the logit function and compared across sites (north, south, and central) and seasons (spring, summer, autumn) using a two-way ANOVA. We ran interactive tests, incorporating the source (stocked vs wild) variable in all analyses as a covariate, along with total length as a scaling factor. A Tukey pairwise comparison was made with mcp() from the multcomp package v1.4-10 in the R statistical environment v3.5.2. (Hothorn et al., 2019; R Core Team, 2018).

## Results

A total of 197 juvenile lake trout (86 wild and 111 stocked, including 15 hatchery-sampled fish) was analyzed for MPTLC. Average (± SD) MPTLC content was 15.2 ± 7.1% of dry mass for stocked fish in the lake and 17.0 ± 6.8% for wild fish. MPTLC of lake trout from the hatchery was 35.1 ± 2.9% of dry mass.

We found no differences in MPTLC among the three Main Lake sites (F_2,175_ = 1.493, p = 0.178) (Figure 2). However, we did find significant differences in MPTLC between stocked and wild fish (F_1,175_ = 27.552, p < 0.001) (Figure 2). In the central and southern Main Lake, wild fish showed significantly greater MPTLC than their stocked counterparts (t_175_ = 3.444, p = 0.008 and t_175_ = 3.438, p = 0.008, respectively).

**Figure 2:**
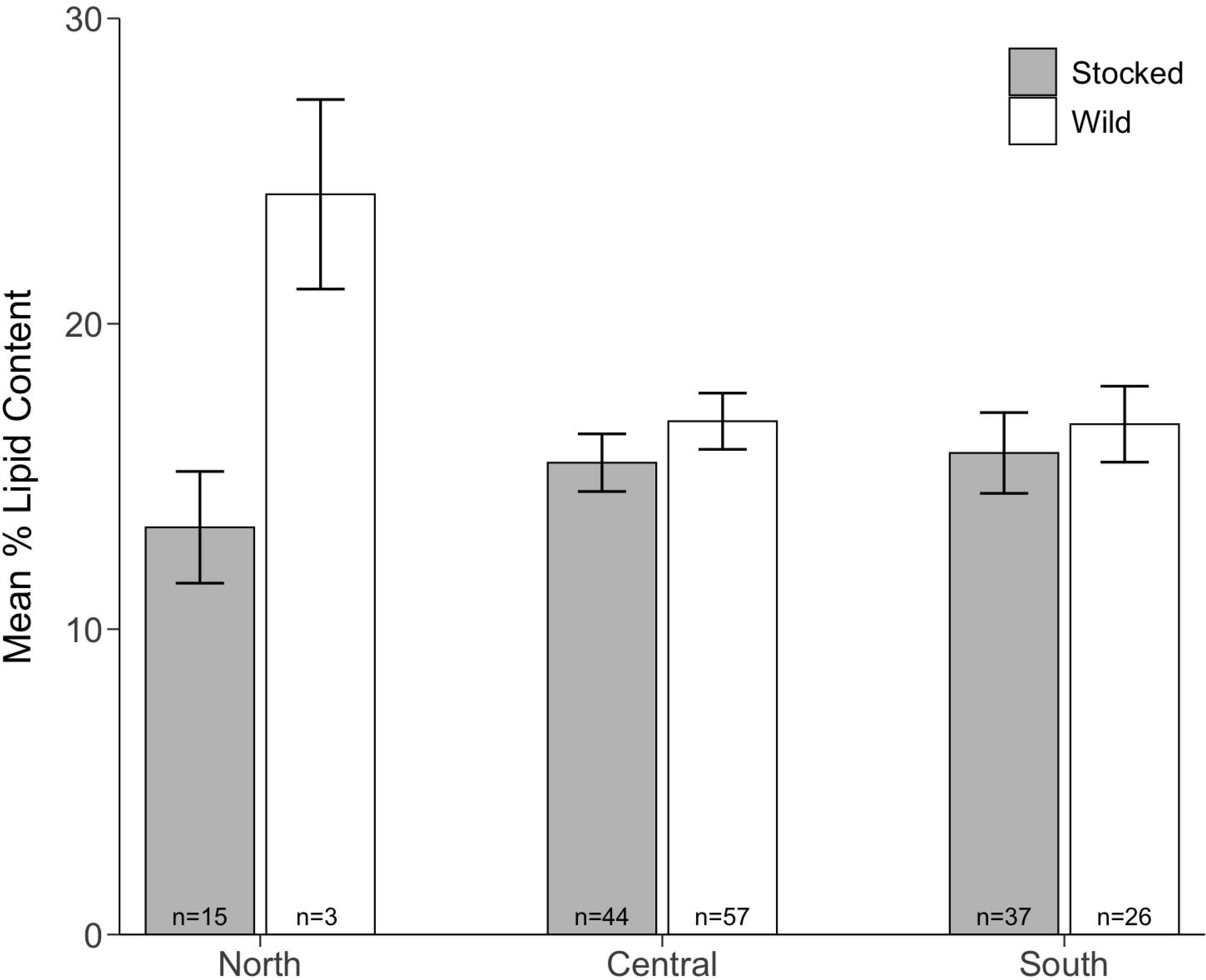
Spatial comparison of mean percent total lipid content of the dry weight of juvenile lake trout ages 1-3 in Lake Champlain captured between 8 June and 29 September 2019. Grey bars denote stocked (fin-clipped) lake trout, and white bars denote wild (unclipped) lake trout. Error bars show standard error. Sample size is indicated at the base of each data bar. North, central, and south refer to the sampling regions in the Main Lake basin.

Juvenile lake trout (wild plus stocked combined) from the central Main Lake varied significantly in MPTLC seasonally (F_2,94_ = 9.858, p < 0.001) (Figure 3). MPTLC was slightly lower in summer (July – August; (t_94_ = −2.702, p = 0.022)) and much lower in Autumn (September; (t_94_ = −4.350, p < 0.001)) than spring (June) for all fish. A pairwise comparison further revealed that stocked fish specifically were lower in mean percent lipid content than wild fish during the summer (t_94_ = 3.209, p = 0.021) and autumn (t_94_ = 2.912, p = 0.049).

**Figure 3:**
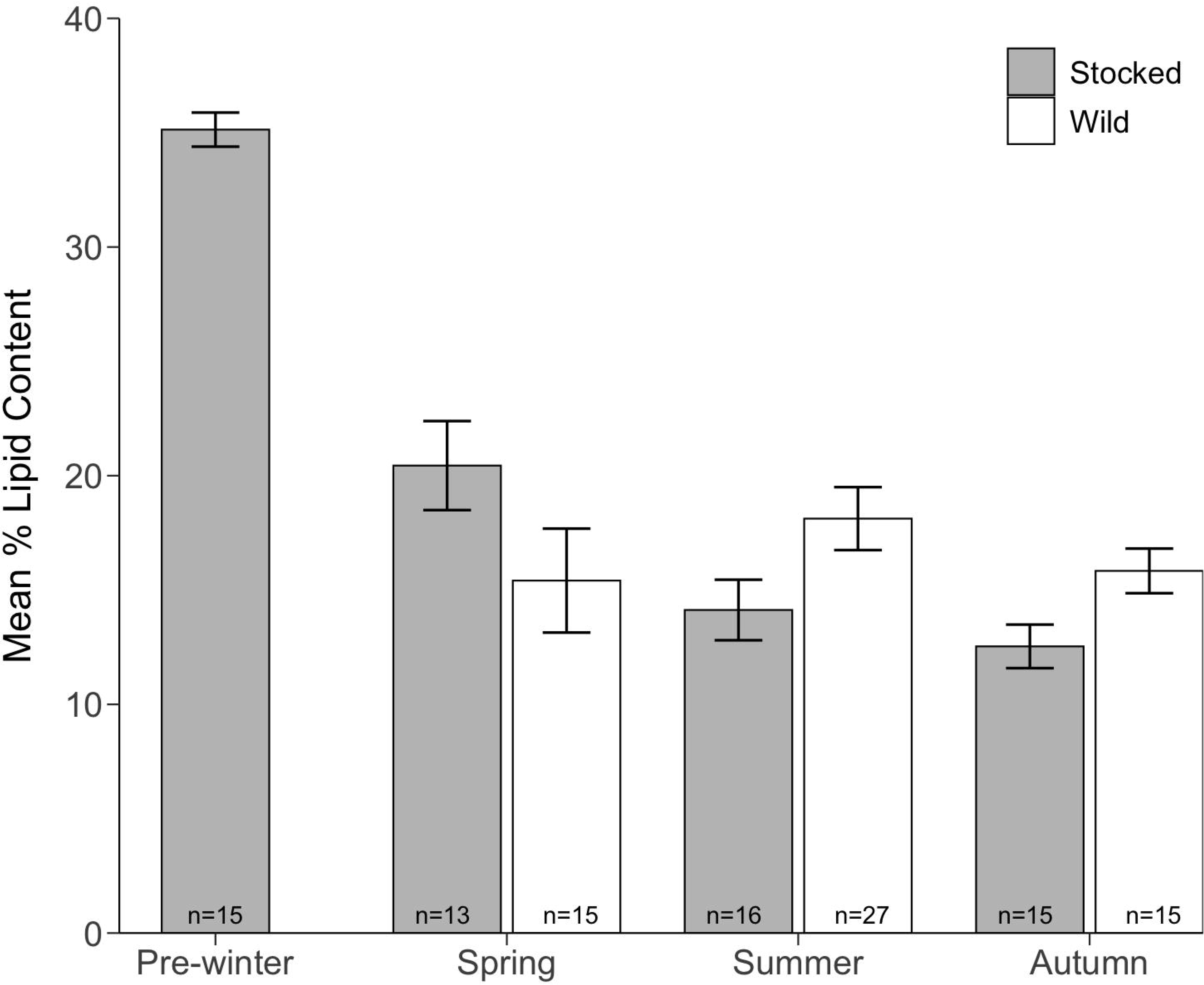
Seasonal comparison of mean percent total lipid content of the dry weight of juvenile lake trout ages 0-3 in Lake Champlain captured between 8 June and 29 September 2019. Grey bars denote stocked (fin-clipped) lake trout, and white bars denote wild (unclipped) lake trout. Error bars show standard error. Sample size is indicated at the base of each data bar. The seasons refer to the month in which lake trout were captured: June (spring), July – August (summer), September (autumn), and a November sample from the Ed Weed Fish Culture Station (pre-winter). We refer to the hatchery sample as pre-winter for comparison between pre-winter and post-winter (i.e., spring) fish.

## Discussion

Our results were unexpected, and each of our hypotheses was refuted. First, we did not find any differences in MPTLC in lake trout sampled from the three different areas of the Main Lake despite higher CPUE and higher proportions of wild lake trout in the Main Lake relative to the northern and southern areas (Wilkins et al. in review). Second, wild lake trout had higher MPTLC than stocked lake trout, despite a two-fold higher MPTLC in stocked lake trout just prior to release into the lake. Further, the high lipid content when hatchery fish were stocked was rapidly lost over their first winter in the lake, and the decline in lipid content continued over summer and into autumn.

We expected that the spatial heterogeneity in abundance of wild juvenile lake trout in Lake Champlain could be due to differences in prey quantity or quality across the different regions of the Main Lake that draw juveniles from the north and south to the central lake. Alternatively, lake trout hatched in the north and south could have lower survival than in the central region if prey resources were higher in the central lake. However, the lack of variation in lipid content among the three regions suggests that lake trout do not experience differences in prey availability across the Main Lake.

Hatchery-reared fish are typically fed a high-ration diet rich in lipids that is reflected in their body composition (Reinitz, 1983). Thus, we expected stocked juvenile lake trout would possess a higher MPTLC than their wild counterparts, similar to other stocked species (e.g., Atlantic salmon *Salmo salar*; Bergstrom, 1989). Analysis of lake trout collected from the Ed Weed Fish Culture Station just prior to stocking showed that hatchery-reared lake trout had a MPTLC approximately two times higher than wild lake trout of the same size in Lake Champlain. However, lipid content of the newly stocked lake trout dropped markedly over their first winter to the level of wild fish and continued to drop throughout summer until by autumn the stocked juvenile lake trout were lower in MPTLC than wild juvenile lake trout of the same age class, although larger in size.

The high lipid content of wild compared to stocked juvenile lake trout suggests that wild lake trout may be more efficient foragers than stocked fish, particularly relative to hatchery fish as they adapt to seeking wild prey. The artificial environment in which stocked fish are raised may not select for traits such as boldness and aggressiveness that are adaptive in natural settings (Brown and Laland, 2002; Brown et al., 2003; Saikkonen et al. 2011). In general, hatchery-raised fish tend to consume less food, fewer prey types, and exhibit reduced ability to switch to new prey types in the wild (e.g., Saikkonen et al., 2011). Density-dependent growth and condition, decreased fin quality, and inferior anaerobic capacity and swim performance have also been documented for fish raised in hatcheries (McDonald et al., 1998). Hatchery-raised brook trout (*Salvelinus fontinalis*) also exhibited lower survival rates once released compared to wild fish because of poor foraging ability (Ersbak and Haase, 1983). The body of evidence suggests that hatchery-raised salmonids are less efficient foragers than wild fish in a natural lake environment, potentially resulting in lower lipid levels compared to wild fish, as we found in this study.

We also found seasonal differences in MPTLC of juvenile lake trout in the central Main Lake. Summer MPTLC was slightly lower than spring lipid levels, and autumn lipid levels were much lower than spring. Our findings contradict patterns reported for other piscivorous fish, where lipids are usually low in the springtime after overwinter depletion, greatest in midsummer months when feeding opportunities are best, and plateau by autumn when system productivity drops before winter (e.g. Madenjian et al., 2000; Metcalfe et al., 2002).

Stocked and wild fish showed different trends in seasonal lipid levels; the pattern in lipid content in the stocked fish appeared to influence the overall trend when all fish were analyzed together. Lipid content of wild fish was consistent with other salmonid fishes, in which lipids are greatest in the summer and lower in spring and autumn (e.g. Madenjian et al., 2000; Metcalfe et al., 2002). In summer, age-1 to 3 lake trout have access to young-of-year smelt and alewife that hatch in June and July, respectively (Simonin et al 2016), and this prey base appears to be sufficient to allow accumulation of lipid storage in addition to growth. Stocked fish, in contrast, showed significant declines in lipid content from spring to summer to autumn. Lipid levels of the hatchery fish also declined substantially between stocking in November and when they were caught in spring, and decreased further through the summer into August. Although this comparison was made between two cohorts (i.e., lake trout sampled prior to stocking in November, and the previous cohort sampled in spring and summer of the same year), hatchery conditions and diet are consistent from year to year, and we can assume reasonable consistency in lake conditions in two consecutive years. The consistent seasonal decline in lipid content of stocked juvenile lake trout suggests that these fish will have less energy reserves than wild juveniles to survive through their second winter in the lake. The high-nutrient diet that stocked lake trout were fed in the hatchery does not appear to give them a lasting advantage over wild lake trout, as wild fish surpass stocked fish in lipid content by the summer following their first winter in the lake. However, the high lipid content of stocked fish may be necessary for survival through the first post-stocking winter, as they learn to feed on active prey and cope with stresses associated with predators.

Data on lipid content can improve understanding of lake trout recruitment in Lake Champlain, inform stocking and conservation efforts, and support the goal of naturally reproducing fish populations. Spatial differences can provide insight on the potential suitability of different areas of the lake to support juvenile lake trout growth, and seasonal differences can provide insight on how fish respond to winter conditions, which may impact juvenile survival rates. The lack of spatial variation in lipid content suggests that the greater abundance of wild recruits in the central Main Lake is not a result of higher feeding. Larger sample sizes and additional years of data would be useful to confirm this result. The increase in lipid levels of wild recruits during the summer is predictable and encouraging, as the data confirm that wild juvenile lake trout are feeding well and therefore have high survival potential. However, we only examined juveniles from June to September. Analysis of juvenile lake trout throughout the year would provide a more complete picture of lipid acquisition and depletion over the winter. The dramatic loss of the lipid advantage of the hatchery lake trout have at stocking is interesting; hatchery fish may be at a substantial disadvantage during their first winter as they acclimate to wild conditions and therefore they need the higher lipid content provided by the hatchery. However, we do not know the survival rate of stocked lake trout during the first winter after stocking; the current survival rate at the current high lipid content supports maintenance of an abundant population, but may not be dependent on high lipid content. If survival is low, hatcheries would benefit from evaluating whether survival could be improved by altering feeding or rearing regimes. We propose two competing hypotheses: high lipid content either 1) provides the necessary energy reserves for stocked fish to acclimate to life in the wild and learn to forage, or 2) imposes a metabolic burden that cannot be sustained in the wild, and reduces the ability of stocked fish to effectively secure necessary energy reserves from a wild prey base. To test these hypotheses, hatcheries could evaluate post-stocking performance and survival of lake trout raised with normal and reduced hatchery diets. If the second hypothesis is supported and the first refuted, hatcheries may be able to rear and stock fewer lake trout with lower ration and maintenance costs to achieve the same survival level.

## Acknowledgements

We thank Pascal Wilkins, Matthew Fidler, and Katrina Rokosz for their tireless support during fish collection, and Robin Sorrentino, Grace Ireland, Samuel McClellan, and Stephen Rotella for their assistance in the laboratory. We acknowledge the University of Vermont Office of Fellowships, Opportunities, and Undergraduate Research for a Summer Undergraduate Research Fellowship, Mini Grant, and Travel Award to MGS.

